# Temporal recurrence as a general mechanism to explain neural responses in the auditory system

**DOI:** 10.1101/2025.01.08.631909

**Authors:** Ulysse Rançon, Timothée Masquelier, Benoit R. Cottereau

## Abstract

Computational models of neural processing in the auditory cortex usually ignore that neurons have an internal memory: they characterize their responses from simple convolutions with a finite temporal window of arbitrary duration. To circumvent this limitation, we propose here a new, simple and fully recurrent neural network (RNN) architecture incorporating cutting-edge computational blocks from the deep learning community and constituting the first attempt to model auditory responses with deep RNNs. We evaluated the ability of this approach to fit neural responses from 8 publicly available datasets, spanning 3 animal species and 6 auditory brain areas, representing the largest compilation of this kind. Our recurrent models significantly out-perform previous methods and a new Transformer-based architecture of our design on this task, suggesting that temporal recurrence is the key to explain auditory responses. Finally, we developed a novel interpretation technique to reverse-engineer any pretrained model, regardless of their stateful or stateless nature. Largely inspired by works from explainable artificial intelligence (xAI), our method suggests that auditory neurons have much longer memory (several seconds) than indicated by current STRF techniques. Together, these results highly motivate the use of deep RNNs within computational models of sensory neurons, as protean building blocks capable of assuming any function.

## Introduction

The activity of a neuron at a given time instant not only depends on its current synaptic inputs but also on its past ones, as well as on a history of its own state. This property is widely taken into account in foundation theories of neural computation (Hodgkin and Huxley (1952), Gerstner and Kistler (2002), Izhikevich (2003)) and extensively observed in the sensory systems of numerous animal species and notably along the auditory pathway where adaptive neural mechanisms such as stimulus-specific adaptation (SSA) (Ulanovsky et al. (2003), Ulanovsky et al. (2004)), short-term plasticity (STP) (David et al. (2007), David and Shamma (2013)), contrast gain control (CGC) (Dean et al. (2005), Dean et al. (2008), Rabinowitz et al. (2011)), context dependence (Asari and Zador (2009), Eggermont (2011)) or priming (Wiggs and Martin (1998), Henson (2003)) were described by experimentalists over the last two decades. Although it is now widely accepted that sound processing in biological systems is constantly modulated by internal representations of recent stimulus statistics (Hershenhoren et al. (2014), Angeloni and Geffen (2018)), the most common computational models of neural processing in the auditory system do not fully reflect these adaptive mechanisms. Indeed, these models are based on spectro-temporal receptive fields (STRFs, Aertsen and Johannesma (1981), Theunissen et al. (2000))) and consist in a cascade of convolutions. Despite their simplicity of implementation and interpretation, they face several issues for fitting neural responses. First of all, their temporal window duration is arbitrary chosen, often without a precise knowledge of integration timescales in the targeted area and neuron class for the given task. Most models actually restrict their STRFs to durations of a few hundred milliseconds (Theunissen et al. (2001), David et al. (2007), Rabinowitz et al. (2012), Thorson et al. (2015), Harper et al. (2016)) whereas numerous electrophysiological studies reported that auditory neurons might integrate their inputs over timescales of a few seconds (Ulanovsky et al. (2004), Asari and Zador (2009)). Secondly, the finite duration of this window creates a sharp boundary between the part of the auditory stimulus that contributes to the response and the one that does not, which is not biologically plausible. Finally, several studies showed that STRF-based models of neural responses in auditory cortex should integrate stimulus information over large temporal windows (hundreds of ms) to improve their neural fitting performances (Machens et al., 2004), despite the fact that such delays are unrealistic in the nervous system (Rahman et al., 2019).

As a partial fix to these issues, conventional models have been extended with nonlinear recurrences akin to the mechanisms mentioned above (SSA, STP, CGC,…), which successfully boost their memory capacity and performances (David et al. (2007), David and Shamma (2013), Willmore et al. (2016), Rançon et al. (2024)), while only requiring a narrow range of delays (Rahman et al. (2019)). These operations are nonetheless only monofunctional add-ons to existing stateless backbones. They still rely on spectro-temporal convolutions and thus keep the problems related to the fixed duration of the temporal window used to process the auditory inputs. Furthermore, most of these studies proposed a different form of recurrence and never attempted to combine them into a single and unified framework, as if they were incompatible, despite the fact that properties such as STP, SSA, or CGC (among others) could be the manifestation of the same and more general computation principle (see e.g., Natan et al. (2015) or Pennington and David (2020)). The manual derivation of the nonlinearities is yet another obstacle to get closer to the underlying mechanism governing neural processing and allowing to better fit sensory responses (Keshishian et al. (2020), Pennington and David (2023)).

To fully address these limitations, we propose here a new class of computational models based on deep recurrent neural networks (RNNs) to characterize sound processing in the auditory pathway. We name these models *StateNets* because they are stateful in time and can potentially internalize any temporal dynamics relevant to reproduce sensory neural behavior (Karmarkar and Buonomano (2007), Barak (2017), Durstewitz et al. (2023)). In line with the observations reported above, their main principle is that the current activity of a neuron relies on an internal and high-dimensional hidden state whose current value depends on previous states as well as on the history of auditory inputs (Buonomano (2000)). To the best of our knowledge, this constitutes the first attempt to model auditory responses with deep recurrent neural networks, despite the fact that recurrent mechanisms have been extensively associated with local microcircuits (Hershenhoren et al. (2014), Costa et al. (2017)) and that audition is a temporal modality. Using electrophysiological data collected in a wide range of animal species and brain areas, we show that StateNet models outperform stateless STRF-based models at fitting auditory neural responses, therefore offering a drastic improvement over the state-of-the-art. In addition, their design is free from unrealistically long synaptic delays and allows to get rid of the crucial hyperparameter that is still manually defined in current modelling studies: the temporal receptive field duration. Furthermore, we also propose a method inspired from AI (Wang et al. (2016), Olah et al. (2017), Selvaraju et al. (2017)) to reveal and interpret STRF-like features for such time-recurrent models, thereby circumventing the lack of explicit spatial-temporal weighting. This method was crafted to closely mimick experimental practices (De Boer and Kuyper (1968), Aertsen and Johannesma (1981), Schwartz et al. (2006)) and generalizes the linear STRF for any model, without any constraint. Our validation benchmark constitutes a precious compilation in the field of auditory neural response modelling, with 11 models (CNNs, a Transformer, and RNNs) tested on 8 datasets, recorded from 3 animal species (ferret, rat, zebra finch) and 5 auditory brain areas (MGB, A1, PEG, MLd, Field L). To foster comparative studies between models, our codes are made publicly available on our online repository (https://github.com/urancon/deepSTRF).

All in all, our study strongly supports the idea that temporal recurrence is a general feature of sensory neurons and demonstrates that combining computational neuroscience and AI opens interesting perspectives for better understanding computations in the nervous system.

## Results

This work presents a new class of models –*StateNet* – based on deep recurrent networks to explain neural responses in auditory system. As observed in electrophysiological recordings, these models have an internal memory and are sensitive to long-term dependencies in their inputs. In the following, we first show that StateNet model performances at neural response fitting tasks are significantly higher than those obtained with state-of-the-art models of auditory processing and also with recent attention-based models (i.e., Transformers). Then, we show that they only need a single timestep to provide accurate temporal predictions, solving the problem of unplausible delays faced by previous models. Finally, we propose a new framework derived from recent developments in explainable AI (xAI) that permits to determine the auditory properties that maximize neural responses in StateNet models.

In the following of this article, we use the terms “RNN” and “stateful models” to designate DNet and StateNet models, in opposition to the remaining models, qualified as “stateless”. Among StateNet models, the GRU, LSTM and Mamba versions are labelled as “Gated RNNs” because of their corresponding recurrent blocks.

### Recurrent networks outperform state-of-the-art models and transformers on a large gamut of datasets

We trained a series of neural network-based models to predict the neural activity of auditory cortical single units, given sound stimuli in the form of spectrograms. We used eight different datasets (AA1 MLd, AA1 Field L, NAT4 PEG, NAT A1, NS1, Wehr, Asari MGB and Asari A1) that combined neural recordings (extra-and intracellular) performed in a large panel of auditory areas and species (rat, ferret and zebra finch). As baseline models, we chose a series of popular backbones, based on the Linear (Nonlinear) Spectro-Temporal Receptive Field (STRFs) model (Aertsen and Johannesma (1981), Chichilnisky (2001), Singh and Theunissen (2003)), as well as a 2D Convolutional Neural Network (CNN) proposed in a recent study and that processes the spectrogram in a similar manner as an image (Pennington and David (2023)). Among these models, DNet (Rahman et al. (2019)) is a recurrent version of the NRF model (Harper et al. (2016)): its hidden units are stateful with leaky dynamics akin to a non-spiking Leaky Integrate-and-Fire (LIF) unit (Gerstner and Kistler (2002)). We also developed a Transformer-based model (Vaswani et al. (2017)) as well as an RNN backbone thereafter referred to as *StateNet* with an interchangeable stateful core. StateNet models included different recurrent architectures: Elman RNNs, LSTM and GRU, S4 and Mamba (Elman (1990), Hochreiter and Schmidhuber (1997), Cho et al. (2014), Gu et al. (2022), Gu and Dao (2023)). Model performances (*CC*_*norm*_) are reported in Table 1.

**Table 1.**
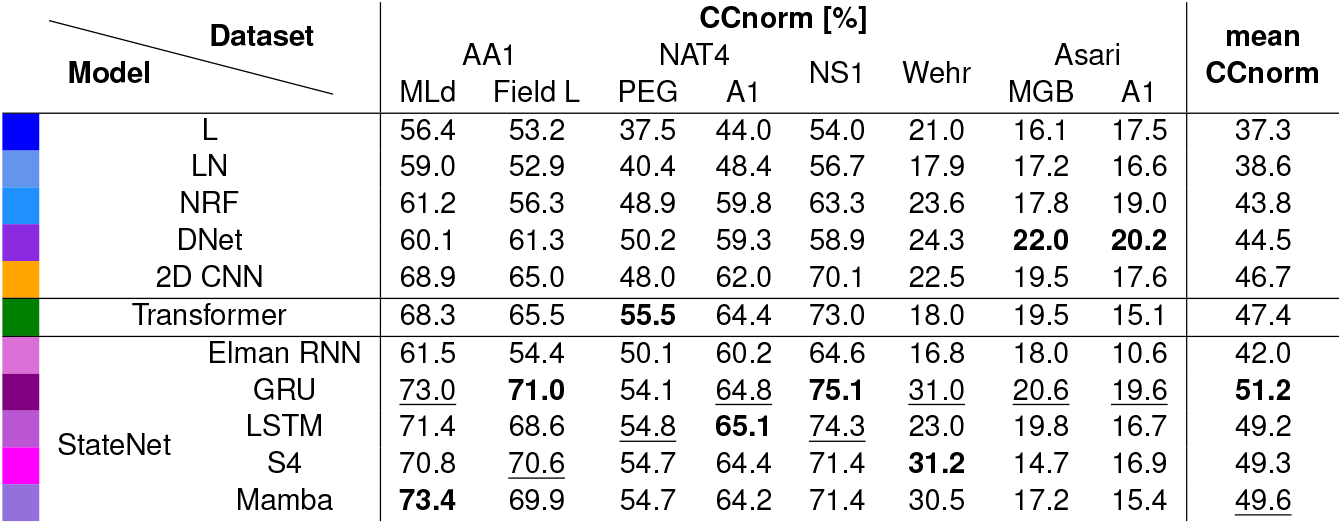
Neural response fitting performances of the 11 tested models for 8 different datasets. Average performances across datasets are provided in the last column. Previous and new (Transformer, StateNet) model classes are respectively shown in the upper and lower rows of the table. Performances are given as the noise-corrected Pearson correlation coefficient, in %. For stateless models, the highest value across the different time windows tested is provided (see Fig. 2). For each dataset and model, *CC*_*norm*_ values were averaged across neurons and training seeds. **Bold** and <u>underlined</u> fonts respectively indicate the best and second best models on a given dataset.

Stateless models were tested using different temporal integration window durations. The best performances across durations are reported here. On average (see the last column in Table 1), except for the simple Elman RNN, all StateNet models significantly outperform the previous state-of-the-art, which is given here by the 2D-CNN models (in yellow, see Pennington and David (2023)). The GRU model notably permits to reach an average *CC*_*norm*_ value of 51.2%, which improves by about 10% the 2D-CNN average score (46.7%). Interestingly, these StateNet models also outperforms Transformers at this task, even though the latter already offers improvement over the previous state-of-the-art on average *CC*_*norm*_ (47.4%). Among StateNet models, the best performances are obtained with the GRU, followed by Mamba, S4, LSTM and finally Elman RNN models. To go further into details, across all datasets, StateNet models are always among the two best ones and lead to the best performances in all datasets except three (NAT4 PEG and Asari MGB and A1). For NAT4 PEG, the LSTM, GRU, S4 and Mamba models outperforms the previous state-of-the-art (∼54.6% on average across the 4 models versus 50.2% for DNet) but provide a lower score than our newly proposed Transformer. For Asari MGB and A1, the best performing model is DNet, which is also recurrent, closely followed by StateNets. To illustrate the better neural fitting performances of StateNet models, we show a few representative examples of model predictions from different datasets in Fig. 1.

**Figure 1.**
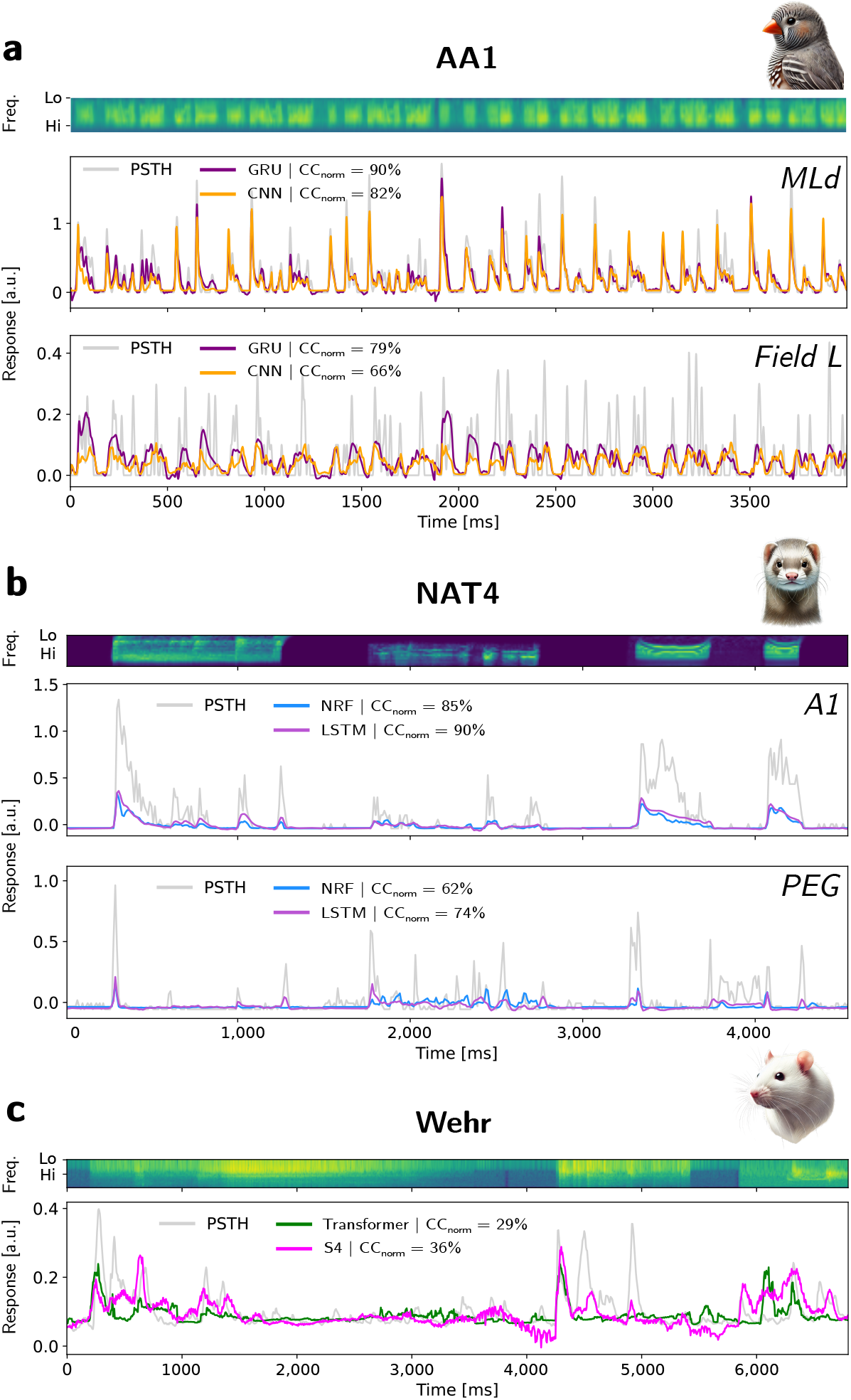
Examples of model predictions vs actual neurophysiological recordings for different datasets. Each panel shows the spectrogram of the stimulus (upper part) and the model predictions (in colors) of the peri-stimulus time histogram (PSTH, in gray) in normalized units (a.u.) as a function of time. *CC*_*norm*_ values are computed over the entire test set of the shown neuron for each model. (**a**) Recordings from neurons in areas MLd and Field L (AA1 dataset, zebra finch). (**b**) Recordings from neurons in areas A1 and PEG (NAT4 dataset, ferret). (**c**) Recordings from a neuron in area A1 and PEG (Wehr dataset, ferret).

### Recurrent networks only need a few timesteps for neural data prediction

Recurrent models (StateNet but also DNet) build their predictions of neural activity at the current time step from a very short temporal window of past stimulus inputs (e.g., 5 timesteps for DNet and as short as 1 timestep for StateNet models). Contrary to previous stateless approaches (L, LN, NRF, 2D-CNN) and to Transformers, these models do not need to process auditory stimuli across time-windows of arbitrary fixed time-durations. To illustrate how performances scale with the duration of the time-window that is accessible, we show in Fig. 2 the performances of the different models as a function of the temporal context. In all datasets (panels a-h), StateNet models are ideally positioned at the upper-left corners of the plots and thus provide an ideal trade-off between performances and temporal context duration. Other models generally increase their performances for longer temporal windows, even though saturation and overfitting effects can be observed in some datasets (e.g., AA1 Md and Field L). Interestingly, the DNet model systematically outperforms its stateless counterpart (NRF model), no matter the length of the input temporal window, and despite an almost equal parameter count. It illustrates the computational advantage brought by implicit recurrence. Furthermore, by learning the adequate temporal integration timescales to fit the data, StateNet models do not depend on the temporal window size hyperparameter, which is otherwise often manually defined to unplausibly long delays (Rahman et al. (2019)).

**Figure 2.**
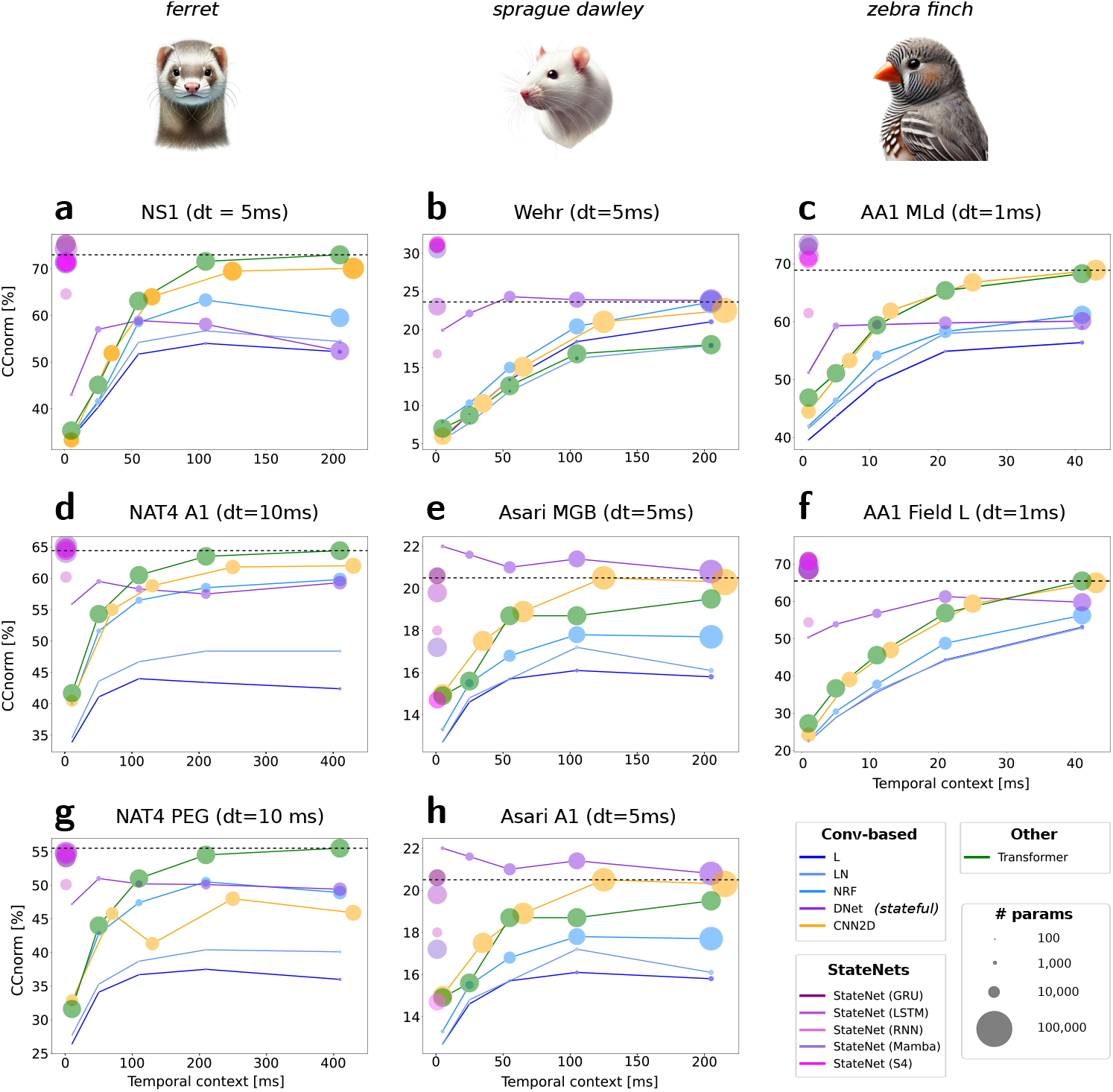
Neural response fitting performances (colored disks) of the 11 tested models for 8 different datasets (panels a to h) as a function of the temporal context (i.e., the duration of the temporal window provided to stateless models in order to predict response in the current time-step). Animal illustrations were AI-generated with DALL-E 3 (OpenAI). The diameters of the disks provide the number of parameters of the associated model.

### A new method to characterize the selectivity and preferred stimuli of auditory neurons in recurrent networks

Auditory selectivity in stateless models can be interpreted by extracting the spectro-temporal receptive fields (STRFs) of neural units (Aertsen and Johannesma (1981), Theunissen et al. (2000)). Such an approach is not directly applicable with StateNet models because they only weight spectral information in an explicit manner while their temporal integration remains implicit. If STRFs can in this case be estimated using reverse correlation techniques based on synthetic (usually white noise, see De Boer and Kuyper (1968) and Aertsen and Johannesma (1981)) but also (although less often) natural stimulus ensembles (see Theunissen et al. (2000), Theunissen et al. (2001)), we propose here a new technique (*“Gradmaps”* and *“Dream STRFs”*) directly inspired from advanced xAI methods (*“Deep Dream”*, see Olah et al. (2017), but also Selvaraju et al. (2017)) and which allows to compute the preferred stimulus of a target unit in any differentiable model (see the “Method” section). A graphic illustration of Dream STRF is provided in Figure 3a. From a null spectrogram (*x*_0_), a forward pass (red arrows) through the frozen network after training provides predicted responses and the associated loss. Then, gradient descent (blue arrows) leads to a gradient map (*g*_0_) which is subsequently subtracted from the initial spectrogram (*x*_0_) to obtain a new stimulus (*x*_1_) that elicits stronger responses in the considered neural unit. This process is repeated until an early stopping criterion is reached, which iteratively leads to spectro-temporal inputs (Dreams) that maximize the neuron’s response.

**Figure 3.**
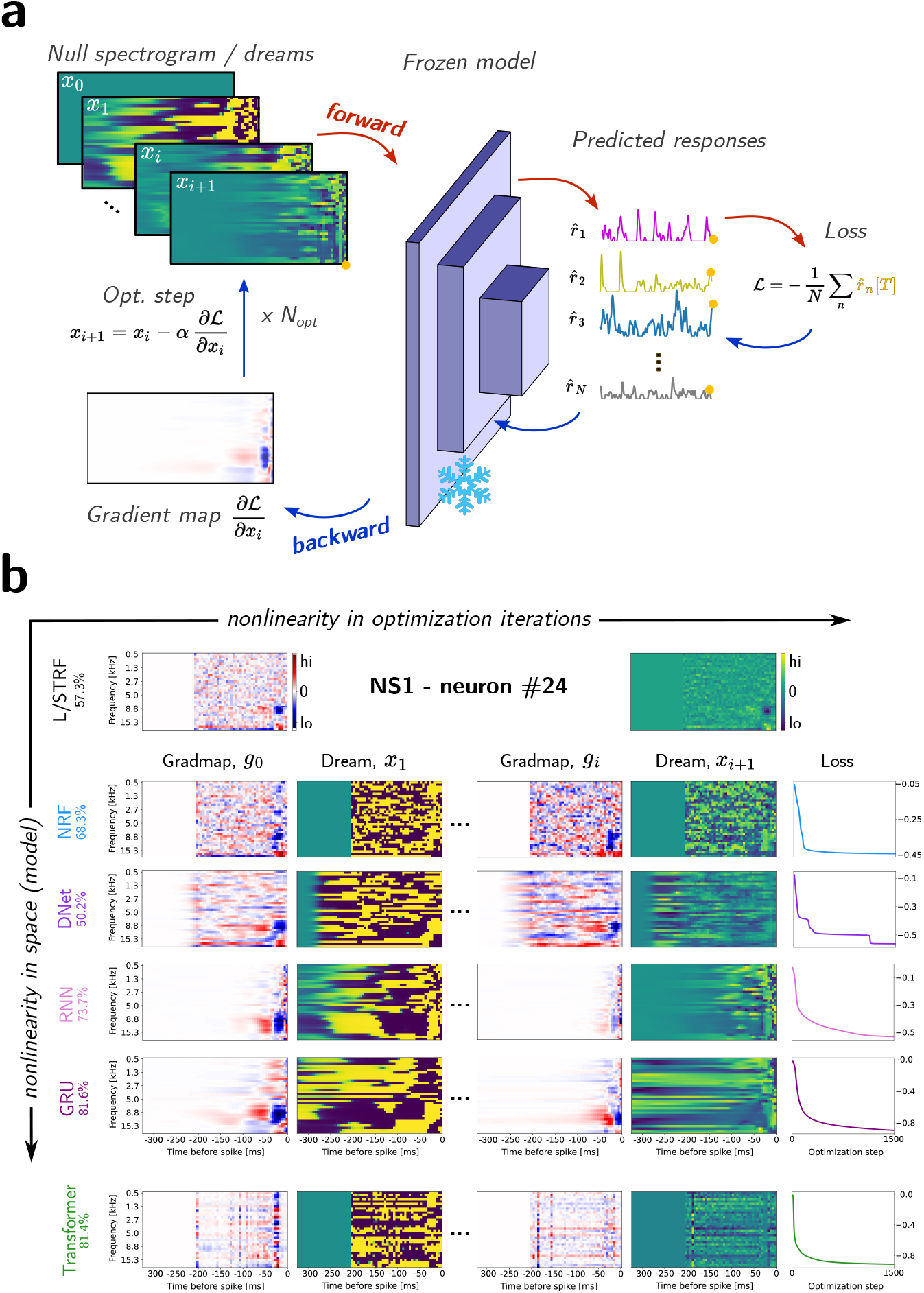
Characterizing the selectivity and preferred stimuli of auditory neurons in StateNet models. (**a**) Illustration of the method. After learning, the parameters of the model are frozen. From a null spectrogram (*x*_0_), a forward pass through the network (red arrows) provides the predicted responses and the associated loss. Then, a gradient descent (blue arrows) leads to a gradient map (*g*_0_) which is subsequently subtracted from the initial spectrogram (*x*_0_) to obtained a new stimulus (*x*_1_) that triggers stronger responses in the considered neural unit. This process is repeated n times and finally leads to the spectro-temporal stimulus (*x*_*n*_) that maximizes the neuron’s response. (**b**) Illustration of the process on one neuron (#24) from the NS1 dataset for different models: NRF, DNet, RNN, GRU and Transformer (*CC*_*norm*_ values obtained with this neuron are displayed with the names of the models). For each model, the panel shows the first gradient map (*g*_0_) and the subsequent Dream (*x*_1_), the last gradient map (*g*_*n*_) and its subsequent Dream (*x*_*n*+1_) and the evolution of the loss function across iterations. For each Dream, we also provide the associated neural responses as a function of time. To improve the readability of the figure, we only show here the 300 last ms before the neuron discharge but for some recurrent models, effects can be observed well before this limit. We provide in Supplementary Fig. S3 the full window of activation obtained with a GRU model for this neuron.

Figure 3b provides examples of gradient maps (GradMaps) and Dream obtained at the initialization and at the end of the process for one single-unit of the NS1 dataset using different models (NRF, DNet, RNN, GRU and Transformer). Other examples obtained from other datasets and models are provided in Supplementary Fig. S3. GradMaps obtained from Stateless (NRF) and stateful (DNet, RNN and GRU) models after the first iteration (*g*_0_) are similar. They show that the targeted neuron tends to be excited (inhibited) by sounds around 8 Hz (15 Hz) presented 50 ms ago. We also observe that Gradmaps of recurrent models are smooth along the temporal axis, which can be explained by the autoregressive nature of the response, one time step being conditioned by the prior. Similarly, smoothness in space/frequency for StateNets can be explained by the initial locally connected (LC) scheme that is applied on this dimension for information compression. Very importantly, Gradmaps also resemble and include the STRF obtained from a linear model (also shown on the figure for comparison). However, GradMaps obtained with StateNet models can extend beyond the temporal window used to compute this STRF. Thus, this technique offers a precise and convenient generalization of the linear STRFs to models of arbitrary depth, width and nonlinearity level. Interestingly, despite their high level of nonlinearity and performance compared to conventional stateless models, the attention mechanism within Transformers leads to much noisier Gradmaps therefore more difficult to relate to STRFs.

Optimized dream STRFs (*x*_*n*_) displayed typical shapes with narrow and wide excitatory and inhibitory regions concentrated in the low latencies. While the energy generally vanishes out at longer latencies for some StateNet models (e.g., RNN), this is not true for all models (e.g., GRU), indicating that some are better than others at leveraging long-term temporal dependencies. Longer latencies are instead highly uniform, indicating that long-past time steps had little influence on the model’s predicted neural response. Typically, dream STRFs effectively span no longer than 50-75 ms. Optimized dream STRFs of conventional stateless models and of the Transformer end abruptly at the limit of their STRF, while they gradually faded away in the case of recurrent models (DNet and StateNet). Interestingly, optimized Dream obtained with the GRU displays complex patterns even in the most ancient time steps, demonstrating the long short-term memory capacities of this model. Importantly, this ability to leverage long-range temporal dependencies is not present in randomly initialized networks, but rather inherited from training on the neural response fitting task (see Supplementary Fig. S3).

One advantage of StateNet models over other approaches is that they produce smooth dreams without the need of regularization. In their case, smoothness directly arises from the temporal recurrence in the models and does not require careful tuning of hyperparameters such as L1 or L2 penalties. A convincing manifestation of this is the smoothness of DNet gradmaps and dreams compared to NRF, which is its stateless counterpart.

In brief, Gradmaps are a generalization of the STRF and represent the linear portion of the neurons’ input-output function. Another way to view them is as the direction towards which the stimulus should change in its space, in order to elicit stronger responses. On the other hand, Dream STRFs are these optimal stimuli that maximize neural activity.

## Discussion

In this paper, we present a new class of models (*StateNet*, see Fig. 4c) based on deep recurrent networks to explain neural responses in auditory system. To the best of our knowledge, we constituted the largest online repository for auditory neural response fitting, with 11 models benchmarked on 8 datasets (AA1 MLd, AA1 Field L, NAT4 PEG, NAT A1, NS1, Wehr, Asari MGB and Asari A1) recorded in large panels of auditory areas (MLd, Field L, PEG, MGB and A1) and species (rat, zebra finch and ferret). We showed that most architectures of this model family (i.e., LSTM, GRU, S4 and Mamba) better fit neural responses than previous approaches proposed in the literature (see Table 1 and also Fig. 2). Notably, the Gated Recurrent Units (GRU) model is the best model on average, and it improves the average *CC*_*norm*_ values obtained by the previous state-of-the-art model (2D-CNN) by about 10%, a considerable amount given the difficulty of the task and the noisiness of auditory neurons.

**Figure 4.**
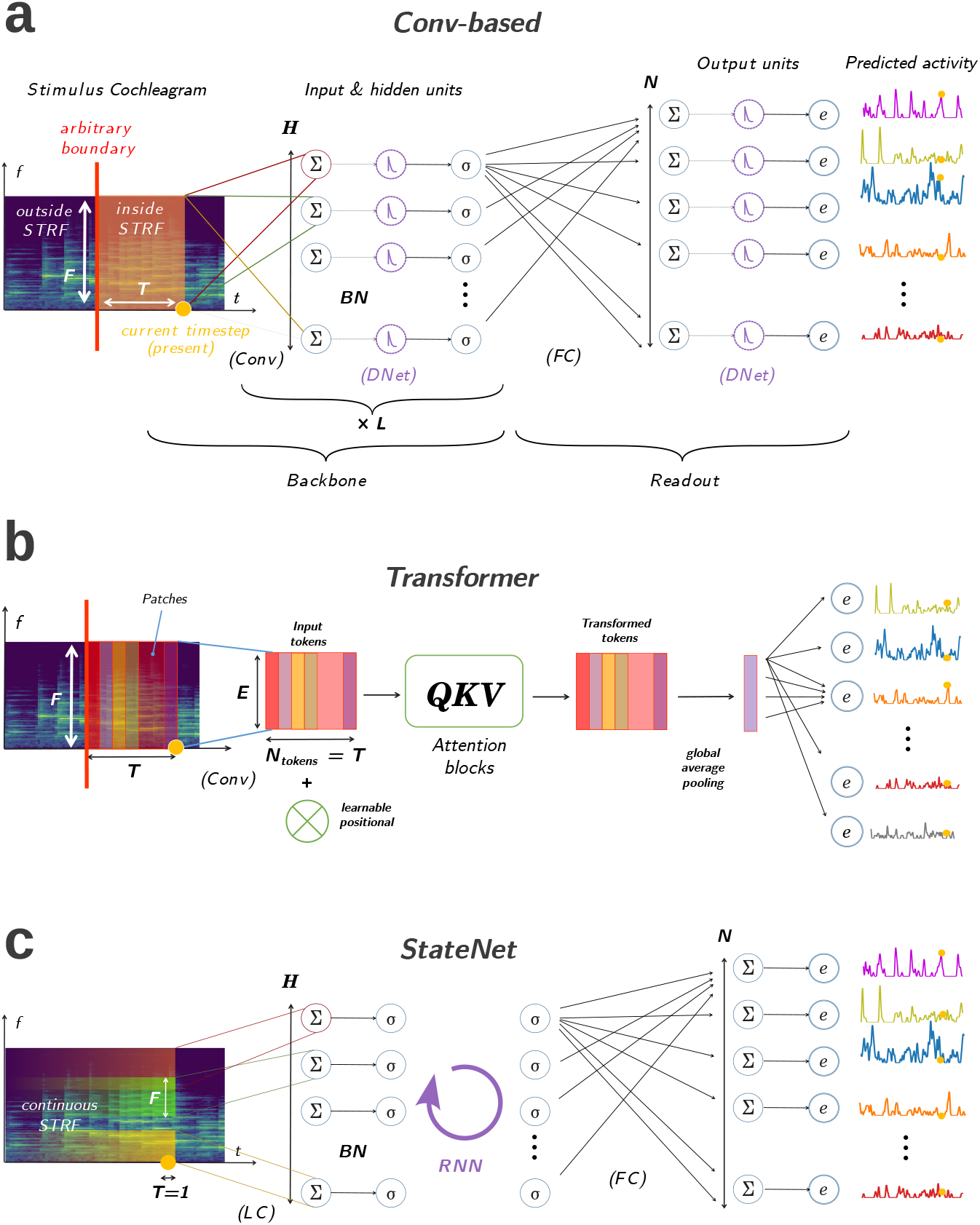
Illustrated architectural features of computational models of auditory neural responses. (**a**) General architecture of canonical conv-based models. The waveform-to-spectrogram transform is not represented here and assumed to have been applied prior to the model. A set of *H* small or large spectro-temporal windows are convolved along the temporal dimension of the cochleagram and followed by some Batch Normalization (BN) and a sigmoid function as an input layer. This operation is repeated *L* times and defines the backbone of the model: learnable parameters and computations that are common to all neuronal units to fit. From the embedding of the stimulus given by the backbone, the readout layer produces predictions of neural activity for each unit in the recorded population, with non-shared, specific parameters including weights, biases and parametric double exponential output nonlinearities. (**b**) Our proposed Transformer model. To give a prediction of current neural activity at time *t*_0_, the network uses a temporal window of size *T* of previous stimulus spectrogram, similar to CNNs. This spectro-temporal window is decomposed into a number of non-overlapping patches which are projected into an embedding space of defined dimension *E* thanks to a 2d convolution. We call the vectors associated to each patch as a result of this operation *tokens*. In our proposed architecture, tokens are the projection of vertical, rectangular patches that are flat, spanning all frequencies but a single time-step. They are then fed to a Transformer block notably composed of several layers of multi-head attention. The output of this block is an equal number of transformed tokens of the same dimensionality, which are averaged into one. From the latter, vector population activity at the current timestep is directly and linearly readout, with a learnable parametric double-exponential output activation function.(**c**) *StateNet* architecture. At each time-step, the current frequency vector is downsampled by a Locally Connected layer maintaining tonotopy, followed by BN. The resulting tonotpic projection is then given to the stateful bottleneck composed of a stateful/recurrent module, which updates its hidden state. Finally, a prediction of neural activity at the current timestep is performed as a linear projection from the latter.

StateNet models (but also the other recurrent alternative, DNet) do not need to process auditory inputs over a time-window of arbitrary fixed duration but instead gradually build their predictions of neural activity at the current timestep from past stimulus inputs (see Fig 3). Thereby, they implicitly learn the extent of their temporal integration window without the need of any extra hyperparameter. It makes them biologically more plausible as durations used in stateless models usually exceed the upper boundary of biological axonal delay distribution (around 50 ms, see Rahman et al. (2019)). Interestingly, most StateNet models also outperform Transformers at fitting auditory neural responses whereas this architecture is now used as a gold-standard in many complex machine learning tasks (Vaswani et al. (2017), Devlin et al. (2019), Dosovitskiy et al. (2021), Zhou et al. (2021)).

If our work is the first to use recurrent deep neural networks to characterize single unit responses in auditory cortex, there were several studies which introduced some form of recurrence into stateless models to capture adaptation mechanisms. To cite a few examples, David and Shamma (2013) proposed a nonlinear stimulus-response model that accounted for synaptic depression. Willmore et al. (2016) used a non-linear pre-processing stage based on a high-pass filtering to replicate adaptation to mean-sound level in the auditory mid-brain. In Rançon et al. (2024), a pair of parameterized temporal filters permitted to capture adaptive mechanisms within ON and OFF responses. All these models led to substantial improvements (although not as pronounced as in the present study) in terms of neural responses fitting performances, thereby reinforcing the idea that recurrent mechanisms are important to characterize auditory responses. Recurrence in these approaches was nonetheless obtained from various add-ons (i.e., non-linearities) to the original STRF-based models and thus did not constitute a core architectural principle, as in the present study.

One possible reason for the better performances of our proposed gated StateNet models is that they can reproduce many forms of adaptive mechanisms described in the literature while stateless approaches (including Transformers) cannot. We provide in the Supplementary materials (see *“Mathematical mapping of RNN networks to adaptation mechanisms observed in the auditory system”*) a mathematical demonstration that gated StateNet models can approximate stimulus-specific adaptation (Ulanovsky et al. (2004)), short-term plasticity (STP, David and Shamma (2013)), contrast gain control (CGC, Dean et al. (2005); Dean et al. (2008); Rabinowitz et al. (2011)) or context dependence (Asari and Zador (2009); Eggermont (2011)). If these computational mechanisms operate at different time scales (David and Shamma (2013)), they might however be complementary (Pennington and David (2020)) and the protean nature of StateNet models offers a unique framework to implement this variety of functions all at once. Recurrent networks are currently the subject of intensive research in the AI community (Beck et al. (2024), Feng et al. (2024)) and we anticipate that in the near future, new models should permit to even better fit neural responses and thereby to improve our understanding of the associated neural mechanisms.

Because StateNet models do not have explicit spectrotemporal weights, classical methods to estimate the auditory selectivity of their neural units are not applicable (Harper et al. (2016), Keshishian et al. (2020)). To circumvent this issue, we proposed here an approach based on advances from the deep learning community (Wang et al. (2016), Olah et al. (2017), Selvaraju et al. (2017)) and which leverages automatic differentiation in recurrent networks. This approach permits for each neural unit to extract its non-linear receptive fields and to estimate the auditory features that maximize its responses (see Fig. 3). It generalizes linear STRFs for any (including nonlinear and recurrent) models, is easy to compute and does not introduce any hyperparameter. It can also be extended to the population level by optimizing the responses of a group of neurons instead of a single unit. One important property of this approach is that it allows for recurrent models to reveal neural selectivity far beyond the temporal windows classically used to estimate STRFs, in line with previous electrophysiological studies which showed that some auditory neurons integrate their inputs over timescales of a few seconds (Ulanovsky et al. (2004), Asari and Zador (2009)). As an illustration, Supplementary figure S3 shows that for many StateNet models, sounds presented more than 2 s before the neuron’s response still modulate the response strength. We believe these extremely large scales of temporal integration to be responsible for the superior explanation power of our deep RNN models.

Our proposed approach is not the first attempt to directly model the nonlinear stimulus-response mappings of auditory cortex neurons with deep neural networks. In a recent work, Keshishian et al. (2020) defined STRFs as the derivative of the output with respect to the input (i.e., the data Jacobian matrix, similar to our GradMaps, see Wang et al. (2016)) in this context. Their method allowed them to find the mathematically equivalent linear function that the neural network applies to each instance of the stimulus, thereby producing linear STRF windows dynamically changing at each time step (DSTRFs) and still restricted to a fixed duration. Moreover, this method assumes strong constraints on the neural network (a feedforward architecture with ReLU activations, no biases in the intermediate layers as well as a linear output) and necessitates to simplify it into an equivalent multilayer perceptron. Our approach does not rely on any temporal window, is totally agnostic on the network architecture (in the present study, it is notably applied on StateNet models but also on more classical models and on Transformers) and thus constitutes a much broader framework to interpret the non-linear receptive fields learned by a model.

Over the last decades, electrophysiologists have estimated the preferred stimulus of a given neural unit using white noise (Aertsen and Johannesma (1981)), synthetic stimuli (Escabi and Schreiner (2002)) or natural sounds (Theunissen et al. (2000)). As the performances of neural encoding models and methods to estimate sensory selectivity progress, it now becomes possible to optimize the features of the stimulus directly during the experiments. Such close-loop approaches have recently been conducted on the visual modality with order-zero optimization methods such as genetic algorithms (Xiao and Kreiman (2020), Wang and Ponce (2022)) but also gradient-based methods (Bashivan et al. (2019), Walker et al. (2019)). Because it is easy to compute and can be applied to any (including recurrent) architecture, the method proposed in the present study could permit to improve these close-loop approaches and to extend them to dynamic stimuli like sounds but also videos. Thus, our work constitutes an important step towards a better characterization and control of neural populations at the unit level, and reaffirms the interest of a hybridization between computational neuroscience and AI on this avenue.

## Materials and Methods

We begin this section with a summary of common models of auditory neural responses and introduce a novel, transformer-based architecture, as well as our fully recurrent model called *StateNet*. Then, we describe the electrophysiology datasets used in this study and set the mathematical framework of the neural response fitting task. Because conventional models and most dataset were already described in a prior study from our group (Rançon et al. (2024)), we invite readers to consult it for a more precise description of conventional models and datasets. Lastly, we explain our process for reverse-engineering models, which generalizes STRFs for arbitrary network depth, width and degree of nonlinearity.

### Canonical computational models of auditory neural responses

The aim of computational models of neural responses in auditory cortex is to convert (“encode”) incoming sound stimuli into time-varying firing rates/probabilities that predict electrophysiological measurements made in auditory areas. Traditionally, these models use the cochleagram of the stimulus –a spectrogram-like representation that mimics processing in the cochlea– and are rate-based (as opposed to *spiking*). Many such models have been proposed in the literature, ranging from simple Linear (“STRF” or “L”) approaches (Aertsen and Johannesma (1981), Theunissen et al. (2000)) to more complex method based on multi-layer CNNs (Pennington and David, 2023). However, all these models use temporal convolutions with finite window lengths, and therefore finite temporal receptive fields (TRFs). In this case, the duration of the TRF is an hyperparameter that is arbitrarily defined by the modelling scientist.

More formally, a model ℳ is a causal application ℝ^*F ×T*^ ↦ ℝ^*N ×T*^ where *F* is the number of frequency bands of a stimulus spectrogram, *T* a variable number of time steps, and *N* a number of units/channels whose activity to predict.

In this paper, we use five models based on this approach: the Linear (L) model (Aertsen and Johannesma (1981), Theunissen et al. (2000)), the Linear-Nonlinear (LN) model (Chichilnisky (2001), Ahrens et al. (2008)), the Network Receptive Field (NRF) model (Harper et al., 2016), the Dynamic Network (DNet) model (Rahman et al., 2019), and finally a deep 2D-CNN model (Pennington and David, 2023). A general architecture of convolutional model is illustrated in Fig. 4a. Because a full description of these approaches was already provided in previous benchmarks (Rançon et al., 2024), we only review here some of their shared general features. We also report minor modifications that we introduced in their implementations so as to make them work on our unified PyTorch pipeline. We invite the reader to consult the original studies that introduced these models for a more detailed description of their functioning.

#### Auditory Periphery

The initial processing stage converts the stimulus sound waveform into a biologically-plausible spectrogram-like representation *x* ∈ ℝ^*F*×*T*^, thereby reflecting operations realized by the cochlea. In the literature, the waveform-to-spectrogram transformation can be performed through a simple short-term Fourier decomposition, or more often through temporal convolutions with a bank of mel or gammatone filters that are scaled logarithmically along the frequency axis. Following the latter, a compressive function such as a cubic root or logarithm is applied. Although the combination of both of these operations make a consensus, there is a variability across studies in their implementation. However, it was shown that such variations are all more or less equivalent and still provide good cochlear sound encodings when modelling higher-order auditory neural responses. As a result, simple transformations should be preferred (Rahman et al., 2020). In order to facilitate present and future comparisons with previous methods, and to limit as much as possible the introduction of biases due to different data pre-processings, we directly use here the cochleagrams provided in each datasets.

#### Core Principle

Classical models rely on a cascade of temporal convolutions with a stride of 1 performed on the cochleagram of the sound stimulus, interleaved with standard nonlinear activation functions (e.g., Sigmoid, LeakyReLU) and followed by a parametric output non-linearity with learnable parameters (e.g., baseline activity, slope saturation value). In all models, the cochleagram is systematically padded to the left (i.e., in the past) with zeroes prior to the temporal convolution operations, in order to respect causality and to output a time series of neural activity with as many time bins as in the input cochleagram.

#### Single unit vs. population fitting

In datasets where all sensory neurons were probed using the same set of stimuli, it is possible for computational models to predict the (vector) activity of the whole population (Pennington and David, 2023). This population coding paradigm allows to train a single model with some learnable parameters shared across all neural units under consideration, and some specific to each unit. As a result, the common backbone tends to learn robust and meaningful embeddings, which further reduces overfitting. Performances are on average better across the population than when fitting an entire model for each unit. Furthermore, this process drastically reduces training time and brings it down to a computational complexity of *O*(1) instead of *O*(*N*), where N is the total number of units in each dataset. For these reasons, we adopt the population coding paradigm whenever possible, that is when various single unit responses were recorded for the same stimuli. This is the case for the NS1, NAT4-A1, NAT4-PEG, AA1-MLd and AA1-Field_L datasets.

#### Output Nonlinearity

All but the L model were equipped with a parametric nonlinear output activation function, learned alongside all other parameters through gradient descent. We used the following 4-parameter double exponential:

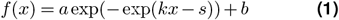

where *b* represents the baseline spike rate, *a* the saturated firing rate, *s* the firing threshold, and *k* the gain (Thorson et al. (2015), Pennington and David (2023)). Importantly, in the case of population models (i.e., when predicting the simultaneous activity of several units, see below), each output neuron learns a different set of these four parameters.

#### Regularization & Parameterization

Canonical models based on convolutions are prone to overfitting and many strategies were proposed to limit this effect, such as the parameterization spectro-temporal convolutional kernels (Willmore and Smyth (2003), Woolley et al. (2006), Ahrens et al. (2008), Thorson et al. (2015). To stick to the most extensively reviewed version of these canonical models, as well as to highlight their limitations, our implementation did not include any such methods. Furthermore, we did not use any data augmentation techniques, weight decay or dropout during training, as it was previously showed that such approaches complexify training and yield little to no improvements in performances (Pennington and David (2023), Rançon et al. (2024)). Instead, we used Batch Normalization (BN), which greatly improved the robustness and performances of all models, including the linear one (L), without compromising its nature as after training, BN’s scale and bias terms can be absorbed by the models own weights and biases.

### Transformer Model

Over the last years, attention-based Transformer architectures (Vaswani et al. (2017)) have been more and more used by the AI community as an alternative to RNN for modeling long sequences (Zhou et al. (2021)), from text (Devlin et al. (2019)) to images (Dosovitskiy et al. (2021)). Contrarily to stateful approaches which rely on backpropagation through time (BPTT) for training, Transformers do not suffer from vanishing or exploding gradients. However, they present some drawbacks, such as quadratic algorithmic complexity scaling with the sequence length. To investigate whether this model is well-suited for fitting dynamic neural responses in auditory cortex, we developed a novel architecture based on the attention mechanism (see Fig. 4b). To the best of our knowledge, it is the first model of its kind that is proposed for this task.

As for stateless models, a hyperparameter *T* defines the length of the temporal context window that serves to predict single unit or population activity at the current time step. Within this window, the spectrogram of the auditory stimulus is projected into *T* tokens (one per time step) of embedding size *E* by means of a fully connected layer applied to each frequency vector. A learnable positional embedding is subsequently added to this compressed spectrogram representation before feeding it to a Transformer encoder with 1 layer, 4 heads, and a dimensionality of 48 (Vaswani et al. (2017)). These hyperparameters were fixed for all datasets. As the outputs of the Transformer encoder are given as *T* processed tokens, we apply global average pooling over the token dimension and use the resulting tensor as the input to a final fully-connected readout layer followed by a double-exponential activation function with per-unit learnable parameters. We observed empirically that the global average pooling operation is crucial to reach good performances while drastically reducing the size of the last fully connected layer.

### *StateNet* Models

A high-level schematic of the processing realized by our *StateNet* models is provided in Figure 4c.

#### Downsampling Locally Connected (LC) layer

At each time step, we downsample the current vector of spectral information to reduce the dimensionality of input stimuli. Contrarily to natural images which are shift-invariant, spectrograms have very different statistics between low and high frequencies, thereby making weight sharing a less efficient computational strategy. Further motivated by the tonotopic organization observed along the auditory pathway (Oliver (2000) or Levy and Reyes (2012)), we use a locally connected (LC) layer with restricted receptive fields (as for convolutional layers) but with independent weights across frequency bands (see Supplementary Fig. S2). In other words, LC contains a subset of the weights of a fully connected (FC) layer, defined by a convolutional (CONV) connectivity pattern over the frequency axis. In theory, the performances obtained with this LC scheme is only a lower bound of what can be reached with FC. However, we found in practice that LC yields overall only slightly lower or similar results using the same hyperparameters (see Supplementary “Ablation study: connectivity in the first layer of StateNet”), but with a smaller number of free learnable parameters, hence reducing the risks of overfitting and permitting a better generalization. LC also outperformed the CONV approach because it relaxes the weight-sharing constraint.

Despite its biological and computational motivations, this approach has only rarely been incorporated into models of auditory processing. For example, Chen et al. (2015) also used a local connectivity for speech recognition, but weight kernels were 2d (spectro-temporal) instead of 1d (only spectral and shared across the temporal dimension). In the field of computational neuroscience, Khatami and Escabí (2020) imposed local Gaussian kernels as fully connected weights. Our implementation differs in that it implements the trade-off between CONV and FC, with fewer parameters than FC, and possibly faster execution.

All in all, our proposed LC downsampling scheme is more biologically plausible than the FC and CONV alternatives, while providing a better trade-off between performances at the neural response fitting task and model complexity. In addition, it executes faster than the FC approach and prior LC implementations. An optimized PyTorch module is available on our code repository.

#### Stateful bottleneck(s)

Because the mathematical details of the modules used here is fully provided in previous studies, we only report below their main properties. We invite the readers to consult the associated papers if a more thorough understanding of their computational principles is needed.

#### Vanilla RNN

*Recurrent Neural Networks (RNN)* are a type of artificial neural networks (ANN) specifically developped to learn and process sequential inputs and notably temporal sequences. They work iteratively and build their output at each timestep from the current inputs as well as a constantly updated internal representation called *hidden state*. In this paper, we designate *“vanilla RNN”* as the classical Elman network (Elman, 1990) natively implemented in PyTorch, often considered as the most naive implementation of this class of models.

#### Gated RNNs: LSTM and GRU

A notorious problem with vanilla RNNs occurs when dealing with long sequences, as gradients can explode or vanish in the unrolledover-time network (Bengio et al. (1994), Pascanu et al. (2013)), preventing them to exploit long-range dependencies and therefore to perform well on large time scales. Gated RNNs such as *Long Short-Term Memory (LSTM)* (Hochreiter and Schmidhuber, 1997) and *Gated Recurrent Units (GRU)* (Cho et al., 2014) successfully circumvent these difficulties, and have imposed themselves as efficient modules for learning sequences in a recurrent approach.

#### State-Space Models (SSMs): S4 and Mamba

*State Space Models (SSM)* is a new class of models that takes inspiration from other fields of applied mathematics, such as signal processing or control theory (Kalman, 1960). Specifically designed for sequence-to-sequence modeling tasks (and thus for time series prediction as in the present study), they build upon the State-Space equations below with various parameterization techniques and numerical optimization methods.

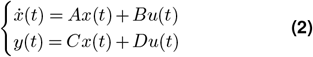

where *u* ∈ ℝ is the input, *x* ∈ ℝ^*N*^ the hidden state vector, *y* ∈ ℝ the output, and *A, B, C* and *D* system matrices. In particular, the original *Structured State-Space Sequence (S4)* model appears as one of the simplest version of this paradigm, with only a few constraints on the system (Gu et al., 2022). At the opposite, the *Mamba* architecture is one of the more recent and sophisticated propositions in which system matrices are input-dependent, and has been shown to perform on par with Transformers on various benchmarks (Gu and Dao, 2023). As a whole, SSMs hold the promise of data-scalable models with great performances, while benefiting from a solid theoretical foundation, which permits to connect them to convolutional models (CNNs), RNNs with discrete timesteps, but also continuous linear time-invariant (LTI) systems of ordinary differentiable equations (ODEs). This last property is particularly interesting as it can ease the reverse-engineering process of a fitted model, thereby allowing for high levels of interpretability. In addition, trained SSMs can easily be modified to work at any temporal resolution, opening interesting use-cases for the field of computational neuroscience and neural engineering.

#### Readout

The readout neural activity for a given unit is computed from a linear projection of the output state vector into a single scalar, repeated at each timestep.

### Electrophysiology datasets of auditory responses

To characterize the ability of models to capture responses in auditory cortex, we fitted them in a supervised manner on a wide gamut of natural audio stimulus-response datasets. These datasets were collected in different species (ferret, rat, zebra finch) and brain areas (MGB, AAF, A1, PEG, MLd, Field L), under varying behavioral conditions (awake, anesthetized) and using different recording modalities (spikes, membrane potentials). All of them are freely accessible on online repositories and were used with respect to their original license. Because the “*NS1*” (Harper et al., 2016), “*NAT4*” (Pennington and David (2022), Pennington and David (2023)) and “*Wehr* “ (Asari et al. (2009), Machens et al. (2004)) datasets have already been used in a previous study from our group and their preprocessing pipelines have been extensively described in the associated article (Rançon et al., 2024), we only describe here the new datasets added to the current study. The corresponding data pre-processing methods are representative of what was performed in the previous datasets.

#### AAA1 datasets: MLd, Field L (zebra finch)

These two datasets consist in single-unit responses recorded from two auditory areas (MLd and Field L) in anesthetized male zebra finches, by Frederic Theunissen’s group at UC Berkeley (Theunissen et al. (2009), Singh and Theunissen (2003), Woolley et al. (2005)). Stimuli were composed of short clips (<10 s) of conspecific songs to which animals had no prior exposure, and were modelled by log-compressed mel spectrograms with 32 frequencies ranging 0 to 16 kHz, at a temporal resolution of 1 or 5 ms. Extracellular recordings yielded a total of 50 single units in each area after spike sorting. Spike trains were binned in non-overlapping windows of 1 or 5 ms matching the resolution of the stimulus spectrogram. Sounds were presented with an average of 10 trials, and the PSTH were obtained for each neuron and stimulus after averaging spike trains across trials and smoothing with a 21 ms Hanning window (Hsu et al. (2004), Willmore et al. (2016)). For each neuron, recordings were performed in response to the same 20 audio stimuli, thereby allowing the training of a single model to predict the simultaneous activity of the whole population (i.e., a “*population coding*” paradigm).

These data, also referred to as “*AA1*”, are freely accessible from the CRCNS website^1^ and were used with respect to their original license.

#### Asari dataset: A1, MGB (rat)

This dataset consists in single-unit responses recorded from primary auditory cortex (A1) and medial geniculate body (MGB) neurons in anesthetized rats, by Anthony Zador’s group at Cold Spring Harbor Laboratory (Asari et al. (2009), Asari and Zador (2009)). Stimuli were natural sounds typically lasting around 2-7 s, and originally sampled at 44.1 kHz and then resampled at 97 kHz for presentation. Their associated cochleagrams were obtained using a short-term Fourier transform with 54 logarithmically distributed spectral bands from 0.1 to 45 kHz, whose outputs were subsequently passed through a logarithmic compressive activation. The temporal resolution of stimulus cochleagrams and responses was set to 5 ms. Because for each cell, recordings were performed in response to a different set of probe sounds, we could not apply the population coding paradigm and one full model was fitted on the stimulus-response pairs for each unit. Contrary to the other datasets, recordings here are intracellular membrane potentials obtained through whole-cell patch-clamp techniques. One remarkable feature of these data is their very high trialto-trial response reliability, making the noise-corrected normalized correlation coefficients of model predictions almost equal to the raw correlation coefficients (see “Performance metrics” subsubection).

Despite the good signal-to-noise ratio of this dataset, some trials are subject to recording artifacts and notably to drifts, which may be caused by motion of the animal and/or of the recording electrode, or electromagnetic interferences with nearby devices. Note that drifts do not contaminate supra-threshold signals resulting from spike-sorted activity such as PSTHs because they are strictly positive and thus have a guaranteed stationarity. In order to remove these drifts, we detrended all responses using a custom approach further explicited in Supplementary section *“Detrending with MedGauss filter”*.

Similar to the previous dataset, these data can be found on CRCNS website^2^ as a subset of the “*AC1*” dataset.

### Neural Response Fitting Task

Because the current benchmark directly builds upon a previous study conducted by our group (Rançon et al. (2024)), performances and model trainings were conducted according to the same methods, which we describe again below.

#### Task definition

Neural response fitting is a sequence-to-sequence, time series regression task, taking a spectrogram representation *x* ∈ ℝ^*F ×T*^ of a sound stimulus as an input, and outputting several 1d time series of neural response (one for each unit), 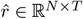. The loss function is the mean squared error (MSE) between the predicted time series and the actual neural recordings.

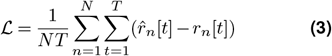

where 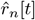 is the predicted neural response for neuron *n* at time-step *t, r*_*n*_[*t*] the corresponding neural recording, *N* is the total number of recorded neurons to fit, and *T* is the total number of time-steps in the time series (to simplify the notations, we drop the time dependencies symbols [*t*] thereafter).

#### Performance metrics

The neural response fitting ability of the different models is estimated using the raw correlation coefficient (Pearsons’ *r*), noted *CC*_*raw*_, between the model’s predicted activity 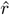 and the ground-truth PSTH *r*, which is the response averaged over all *M* trials *r*^(*m*)^:

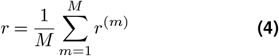

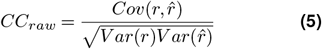

where the covariance and variance are computed along the temporal dimension. Due to noisy signals and limited number of takes, perfect fits (i.e. *CC*_*raw*_ = 1) are impossible to get in practice. In order to give an estimation of the best reachable performance given neuronal and experimental trial-to-trial variability, we use here the normalized correlation coefficient *CC*_*norm*_, as defined in Hsu et al. (2004) and Schoppe et al. (2016). For a given optimization set (e.g., *train, validation* or *test*) composed of multiple clips of stimulus-response pairs, we first create a long sequence by temporally concatenating all clips together. We then evaluate the signal power *SP* in the recorded responses as:

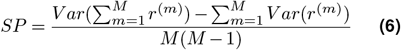

which allows to compute the normalized correlation co-efficient:

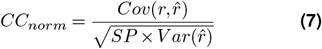

When only one trial is available, we set *CC*_*raw*_ = *CC*_*norm*_, which corresponds to a fully repeatable recording uncontaminated by noise, thereby preventing any overestimation of performances by setting a lower bound in absence of data.

#### Optimization process / model training

All models were trained using gradient descent and backpropagation, with AdamW optimizer (Loshchilov and Hutter (2019)) and its default PyTorch hyperparameters (*β*_1_ = 0.9, *β*_2_ = 0.999). We used a batch size of 1 for all but NAT4 datasets, which have a limited number of training examples, and a batch size of 16 for both NAT4 datasets, which have consequently more. The learning rate was held constant during training and set to a value of 10^−3^. We found empirically that these values led to better results.

We split each dataset into a training, a validation and a test subset respecting a 70-10-20% ratio as much as possible, depending on the number of stimulus-response pairs available for each cell in each dataset. At the completion of each training epoch, models were evaluated on the validation set and if the validation loss had decreased with respect to the previous best model, the new model was saved. Models were trained until there were no improvement during 50 consecutive epochs on the validation set, at which point learning was stopped, the last best-performing model was saved and evaluated on the test set. This procedure was repeated 10 times for different train-valid-test data splits and model parameters initializations, and the test metrics were averaged across splits.

This unified approach for implementing and optimizing the parameters of each of the models (i.e., using the same regularization method, fitting approach, number of cochleogram channels, …) allows a fair comparison between them. Indeed, as all the models (StateNet models but also L, LN, NRF, DNet and 2D-CNN) were constructed using exactly the same pipeline, it implies that a model with higher neural fitting performances is genuinely better. Note that this homogenisation strategy necessarily introduced differences between our general pipeline and those of the studies that originally described these models.

### Model interpretability with feature visualization: gradient maps and deep dreams

We propose here a gradient-based iterative method which, for each unit of a trained neural network ℳ, extracts its non-linear receptive fields and estimates the auditory features *x*_*i*_ that maximize its responses. This method builds on feature visualization techniques originally introduced in the AI community (Wang et al. (2016), Olah et al. (2017), Selvaraju et al. (2017)) is also known as gradient ascent, leveraging the fact that all the mathematical operations in our models are differentiable. The different steps of this approach are illustrated on Fig. 3a and can be summarized as follows:

1. As the first input to the model, use the null stimulus *x*_0_ ∈ ℝ^*F ×T*^, a uniform spectrogram of constant value (0 in our case). This initial stimulus is unbiased and bears no spectro-temporal information. From an information-theoretic perspective, it has no entropy. For a parallel with electrophysiology experiments, it is worthy to notice that the spectrogram of white noise is theoretically uniform too. The absence of spectro-temporal correlations within probe stimuli (which we respect here) is a strong theoretical requirement that led to the use of white noise in the initial development of the linear STRF theory (Aertsen and Johannesma (1981)), while further studies preferring natural stimuli used advanced techniques to correct their structure (Theunissen et al. (2000)).
2. Pass *x*_0_ through the model and compute its outputs (i.e., the predicted time-series of activation for the whole neural population):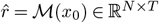
3. Define a loss ℒ: ℝ^*N ×T*^ ↦ ℝ to minimize and compute it from the model prediction 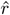. In this paper, we only targeted a single unit *n* and used the opposite of its activation at the last (*T*-th) time-step: 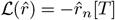. To compute the STRF associated with any neural population 𝒩, the following general loss can be used: 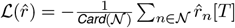. The choice of this loss function permits to make the connection with the Spike-Triggered Average (STA) approach used by electrophysiologists and where the stimulus instances preceding the discharge of the target unit are averaged (De Boer and Kuyper (1968), Aertsen and Johannesma (1981), Schwartz et al. (2006)). Models in our study were fitted on PSTHs or membrane potentials and thus output floating point values, which can be viewed as a spike probability. Trying to maximize this value at the present time-step (the last of the time series) by constructing previous stimulus time steps closely mimics STA. Maximizing the average firing rate across the whole stimulus presentation could be interesting to investigate in future studies.
4. Back-propagate through the network the gradients of this loss 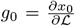, thereafter referred to as ***GradMaps***. From their definition, these GradMaps can be directly related to linear STRF (see Supplementary text *“Bridging the gap between STRFs, gradmaps and dreams: theoretical framework”*).
5. Use these gradients to perform a gradient ascent step and modify the input stimulus. For simplification, we denote this operation as *x*_1_ = *x*_0_ − *αg*_0_, but some optimizers have momentum and more elaborate update rules. This is notably the case of Adam and AdamW (Loshchilov and Hutter (2019)), which were used in our study. We did not use the SGD optimizer as it converged to much higher loss values –so less optimal– in preliminary experiments.
6. Repeat these steps until an early stopping criterion is satisfied, in our case after a fixed number of iterations (1500, a value which led to sufficient loss decreases, see curves in Fig. 3). The result of this process is an optimized input spectrogram *d* = *x*_*i*_ that maximizes the activation of the target unit(s), thereafter referred to as ***Dream***. If we define ℱ: ℝ^*F ×T*^ ↦ ℝ^*F ×T*^ as one iteration of the above process, such that *x*_*i*_ = ℱ (*x*_*i*−1_|ℳ, ℒ, *n*) then recursively get *x*_*i*_ = ℱ ○ ℱ ○ … ○ ℱ(*x*_0_|ℳ, ℒ, *n*) = ℱ^*i*^(*x*_0_|ℳ, ℒ, *n*).

This approach does not assume any requirement on the model to interpret. It can produce infinitely long GradMaps, which are relatable to linear STRFs and can be implemented on any architecture, including RNNs like StateNet but also stateless and transformers.

### Reproducibility

One major contribution of this work is the sheer amount of implemented models and compiled datasets. All models were implemented and trained using PyTorch, a gold-standard library for deep learning in Python, leveraging autodiff. Datasets were pre-processed under the same format of a PyTorch Dataset class for convenience. Upon publication of this article, we will make our code freely available on Github at the following address: https://github.com/urancon/deepSTRF. It will complement the already existing basis of this repository, originally developed in a previous work of our group (Rançon et al., 2024). The aims of this library are to normalize practices in the field of neural response fitting with quantitative computational models, to provide a common ground for benchmarking and to permit a fair comparison between models on various preprocessed datasets, with ready-to-use and user-friendly classes and methods.

Jobs required less than 2 GiB and were executed on Nvidia Titan V GPUs, taking tens of minutes to several hours, depending on the complexity of the model. As an example, the population training (5 seeds) of the StateNet GRU model trained in population coding on all 73 NS1 neurons in parallel typically takes less than 10 mins. Conversely, the single unit training (5 seeds) of the same model on the 21 neurons of the Wehr dataset takes more than 3 hours.

## Detrending raw signals with “MedGauss” filter

We demonstrate here the need of detrending neural signals in Wehr and Asari (CRCNS AC1) datasets, consisting of in-vivo patch clamp recordings, akin to drift and noise. We compare in Fig. S1 below the output of a simple linear detrend to our “*MedGauss*” filtering solution, which aims at minimizing manual curation. In this method, we apply a median low-pass filter (2.5 s temporal window) followed by a gaussian (200 ms of standard deviation) and subtract the output from the raw response:

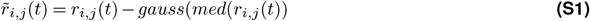

where *r*_*i*,*j*_ represents the *j*th response trial to the *i*th stimulus of a given neuron. The main motivation for this approach is that a single polynomial detrend (as available in common tools such as MATLAB or Scipy) would not be able to correct artifacts of different orders occurring before/after periods of stationarity. This process is illustrated in Supplementary Fig. S1.

**Figure S1.**
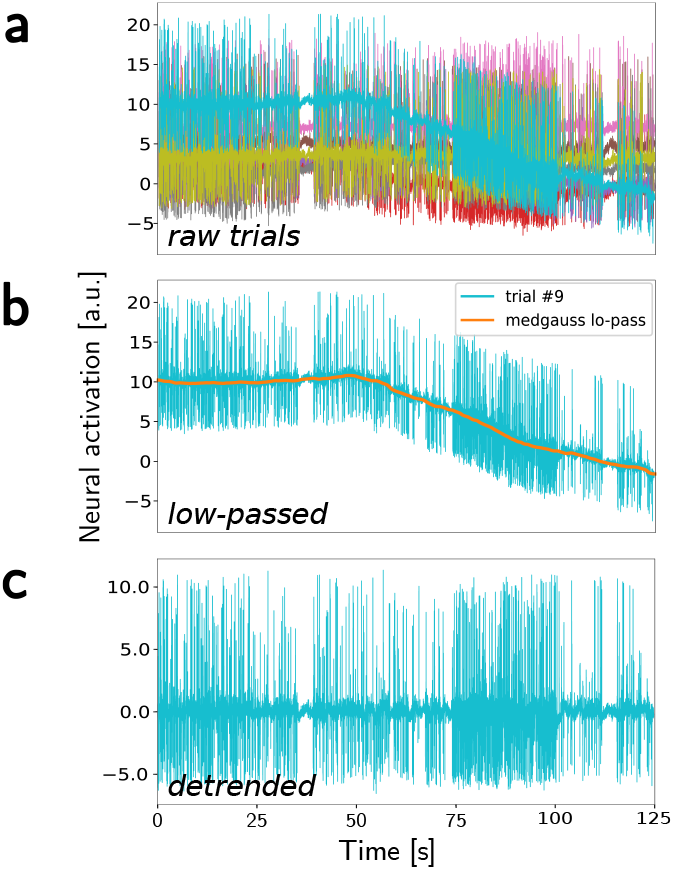
Illustration of the so-called “MedGauss” detrending. (**a**) Response trials of Asari MGB neuron #5 to stimulus #6. They exhibit different temporal means, and some global (i.e. temporally long-range) corruption. (**b**) Response trial of interest and subject to drift, with its low-passed version. (**c**) The detrended signal.

The response trial presented in the above figure shows a spike signal with very nonlinear and long-duration drift. Therefore it cannot simply be detrended with a linear, or even quadratic detrend. Even more generally, polynomial detrend has the inconvenient of requiring the definition of a hyperparameter, the order, which might differ from one recording to another depending on the type of corruption. Our detrending process can correct global and highly nonlinear drifts of very different shapes, therefore removing the need for manual intervention.

## Ablation study: connectivity in the first layer of StateNet

We trained the StateNet GRU model on each dataset with fully connected (FC), Locally Connected (LC) and convolutional (CONV) connectivities as the first layer (see Figure S2). The results are reported below in Table 2.

**Table 2.** Performances of the StateNet GRU model on each dataset with various connectivity schemes for spectral downsampling. Normalized correlation coefficients are given in %, and learnable parameter numbers are given for a backbone with only one output neuron. **Bold font** indicate the best connectivity pattern among the three options available. - symbolizes duplicated entries (see above cell).

| Dataset |  | F | K | S | C | # Params |  |  | CCnorm [%] |  |  |
| --- | --- | --- | --- | --- | --- | --- | --- | --- | --- | --- | --- |
|  |  |  |  |  |  | FC | LC | CONV | FC | LC | CONV |
| NS1 |  | 34 | 7 | 3 | 7 | 32,355 | 30,465 | **29,961** | 73.8 | **75.1** | 74.4 |
| NAT4 | A1 | 18 | 5 | 2 | 10 | 31,241 | 30,331 | **29,971** | **54.7** | 54.1 | 52.5 |
|  | PEG | - | - | - | - | - | - | - | 64.6 | **64.8** | 63.3 |
| AA1 | MLd | 32 | 7 | 3 | 7 | 26,349 | 24,774 | **24,326** | **74.4** | 73.0 | 69.4 |
|  | Field L | - | - | - | - | - | - | - | **71.6** | 71.0 | 69.6 |
| Wehr |  | 49 | 9 | 4 | 5 | 21,296 | 19,096 | **18,596** | **32.3** | 31.0 | 31.6 |
| Asari | A1 | 54 | - | - | - | 25,331 | 22,631 | **22,081** | **23.9** | 19.6 | 23.3 |
|  | MGB | - | - | - | - | - | - | - | 20.2 | 20.6 | **21.7** |

As expected, the number of parameters in StateNet-LC models is higher than StateNet-CONV but lower than StateNet-FC (because LC is only a subset of FC). In terms of performances, LC almost always surpasses CONV, and yields slightly lower *CC*_*norm*_ than FC. Therefore, LC offers an interesting trade-off between neural response fitting accuracy and model size.

**Figure S2.**
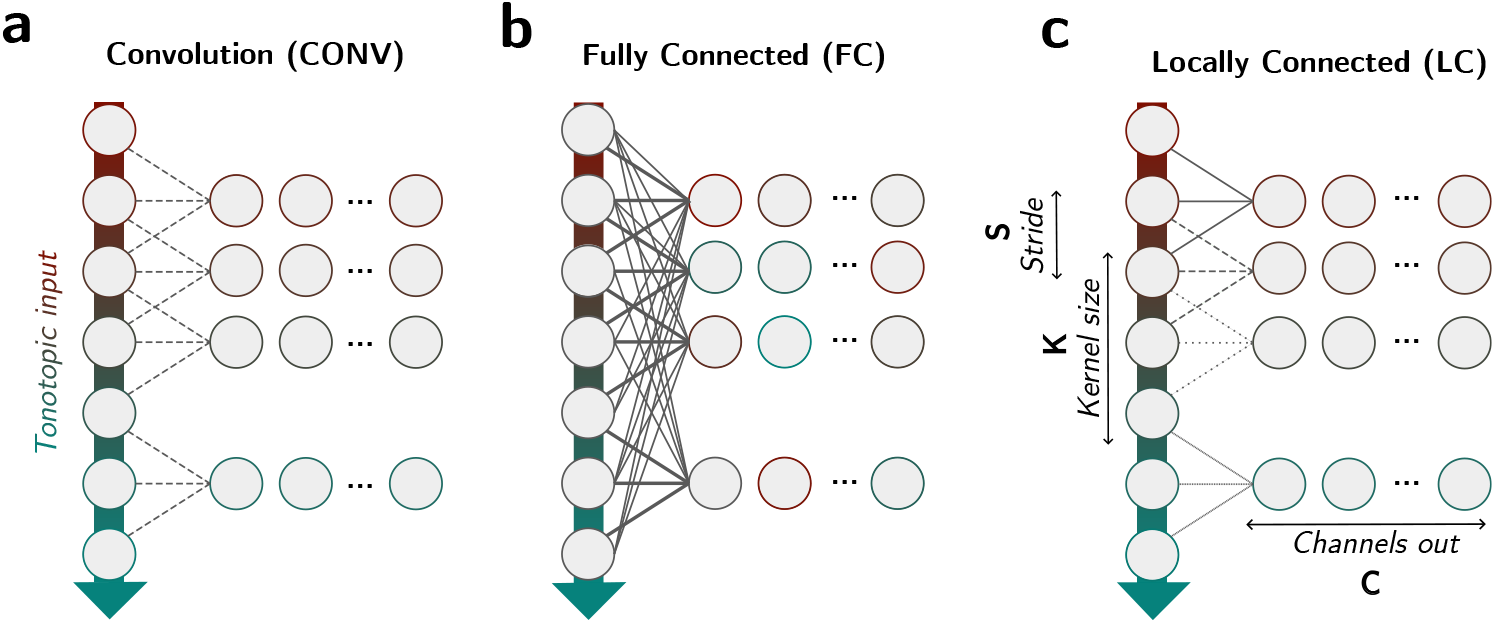
Illustration of the proposed Locally Connected (LC) connectivity compared to Fully Connected (FC) and convolution (CONV). Example proposed for one-dimensional inputs, such as frequency bands, as used in StateNet. In a CONV layer, the same weight kernel is applied at each location of the input, while in an FC layer, each input-output neuron pair has a specific weight. LC is essentially a mix between both: a CONV but without weight sharing, or in other terms, a subset of synaptic connections present in FC.

## Gradmaps and full dreams for other NS1 neurons

In Figure 3, we show dreams of one neuron from the NS1 dataset, truncated in the past to focus on the most recent timesteps, for better readability. As a demonstration of the robustness of the approach, we show in Supplementary Fig. S3 that gradmaps of complex StateNet models generalize conventional STRFs for the other neurons of this dataset. This figure also illustrates the temporal extent of context dependence in auditory neurons, as the preferred stimulus (*“dream”*) for this neuron presents complex patterns several seconds prior current activation in the best-performing StateNet models (GRU, LSTM, S4, Mamba). The complex patterns that have emerged from the optimization procedure in the very long latencies for those models are *not* artifacts: removing them and presenting a truncated version of half the most recent time-steps of their respective dream to those models leads to lesser activations at the current time (right side of the time axis). A possible reason could be that the model needs to be at some unstable point of equilibrium (i.e., in the hidden state space) in order to generate sudden, high activations. Thus, our results suggest that when trying to elicit high responses in auditory neurons, long (several secs) *“warmup”* stimuli are necessary to be presented to the animal. Another marking fact from our qualitative analysis of StateNet dreams, is the repetition of some warmup patterns (e.g., see GRU on Supplementary Fig. S3), despite the fact that each time-frequency bin was optimized independently from the others (neither structural constraint nor regularization was applied). Finally, this capacity to leverage long-range dependencies is not solely enabled by the model architecture, but also inherited from the neural response fitting process, as a randomly initialized GRU (*“randGRU”*) model displays very short dreams (other neurons not shown).

## Bridging the gap between STRFs, gradmaps and dreams: theoretical framework

This section aims at consolidating the theoretical relationship between GradMaps, Dreams, and what is commonly referred to as *“linear STRF”* in the literature. This latter term has been used interchangeably as the neuron’s “preferred input” eliciting maximal activity (Rabinowitz et al. (2012), Keshishian et al. (2020)), or as its linear stimulus-response “transfer function” (Aertsen and Johannesma (1981), Machens et al. (2004), Thorson et al. (2015)). As we will see below, those definitions are related, in particular in the case of the linear (L) model, hence our choice to disentangle “GradMaps” from “Dreams”.

Let ℳ^∗^ be the ideal model of a neuron such that ℳ^∗^(*x*) = *r*, ∀*x ∈* ℝ^*F ×T*^ with ℳ^∗^(*x*) the “true” function implemented by the neuron. Then, the “true” linear STRF of the neuron is defined as the gradmap 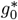 such that:

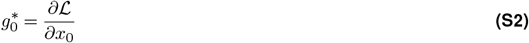

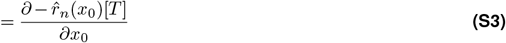

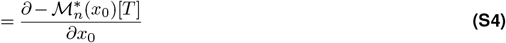

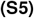

Thus, when a given model ℳ tends towards the optimal model ℳ^∗^ that perfectly captures the function implemented by the neuron on the whole stimulus space, we get:

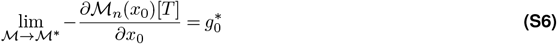

It should be reminded that the *CC*_*norm*_ metric is only a proxy for the distance separating ℳ from ℳ^∗^: it is computed on a subset of stimuli that is limited by experimentation. In any case, as we get better performances at the neural response fitting task with the model ℳ, we can expect experimental STRFs to tend towards the neuron’s “true” STRFs.

From our definitions in the Materials and Methods section, and the fact that *x*_0_ = 0 is the null vector of the stimulus space ℝ^*F ×T*^, the order-1 dream *x*_1_ is:

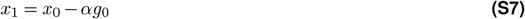

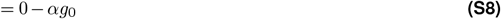

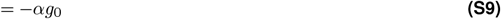

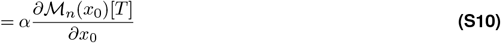

So it turns out that the linear STRF *g*_0_ and the order-1 dream are equal, up to a minus sign and a scaling constant. The particular case of ℳ being the Linear model further fueled non-rigorous definitions for the STRF. The L model can be simply written as ℳ = *Wx* + *b*, with *W* and *b* as the weight matrix and the bias vector. Updating the Eq. S10 yields:

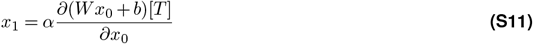

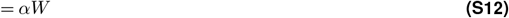

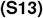

Then, generalizing to any other dream optimization step *i*:

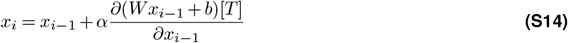

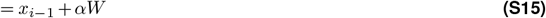

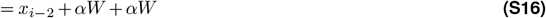

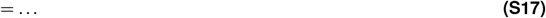

It recursively follows that for the L model:

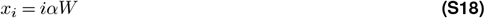

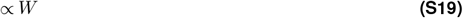

We now better understand why the term *“STRF”* has been used to designate both the preferred stimulus of a neuron, as well as changes in stimuli leading to changes in activity (Rabinowitz et al. (2012)), or as its convolutional kernel (transfer function): this essentially stems from the linear assumption, and vanishes in our framework, in particular in nonlinear regimes.

## Mathematical mapping of RNN networks to adaptation mechanisms observed in the auditory system

In this paragraph, we demonstrate how deep RNNs can approximate adaptation mechanisms observed in the auditory cortex. We first focus on the gated Recurrent Unit (GRU) because this model leads to the best performances at the neural response fitting task (see Table 1), and then consider other RNN models.

**Disclaimer 1**: If the nonlinearities in the GRU model prevent it from strictly implementing some adaptation mechanisms, it can still approximate them. In the case where a strict mathematical equivalence is required, working in the linear regime of the *σ* and `tanh` functions can lead to better approximations.

**Disclaimer 2**: If the demonstrations provided here show that the GRU model can approximate these mechanisms, it does not imply that it is precisely what it learned during the training process. In practice, we can expect that the model learns a mixture of different mechanisms.

### Reminder on GRUs equation

The equations of the Gated Recurrent Unit (GRU) are the following:

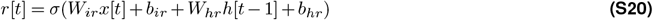

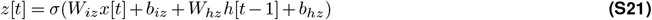

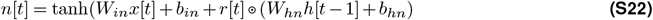

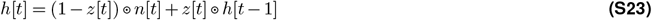

where ⊙ denotes the element-wise vector product (Hadamard product).

#### Leaky integration with GRUs

The simplest and most widespread model of neuronal activity is the Leaky-Integrate and Fire (LIF), representing the electric potential of a neuronal membrane viewed as a RC circuit:

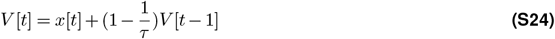

This equation is reminiscent of Eq. S4, when *V* = *h* and 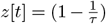. Doing so, 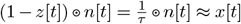 in Eq. S4 with *W*_*in*_ = *τ*, *b*_*in*_ = 0, *r*[*t*] = 0, *W*_*hn*_ = 0 and *b*_*hn*_ = 0. Hence, GRUs can approximate models with an exponential leaky memory, such as the LIF.

### Non-leaky integration

GRUs can also implement non-leaky integration, when *z*[*t*] = 1 ∀*t*, e.g. *W*_*iz*_ = *W*_*hz*_ = 0 and *b*_*iz*_ or *b*_*hz*_ ≫ 0. It allows the model to remember input patterns over very long periods, and to modulate current responses based on a long stimulus history. Because of their vector formulation, some elements of the hidden state can be retained with little or no leak, while others can be more easily replaced by new features. For instance, the first element *h*_1_ of the hidden state can correspond to a permanent memory of some input pattern if *z*_1_[*t*] = 1, while *h*_2_ can be constantly updating if *z*_2_[*t*] = 0.2.

Furthermore, we can construct a GRU model where *h*_1_ has a non-leaky memory (i.e., *z*_1_[*t*] = 1) for most of the inputs, but is still capable of important updates (i.e., *z*_1_[*t*] ≈ 0) when encountering specific patterns. This can be achieved by letting *z*_1_[*t*] depend on *x*[*t*] and *h*[*t* − 1] through non-zero *W*_*iz*_ and *W*_*hz*_ matrices, as opposed to the previous examples where it was set to a constant leakage rate (related to the time constant *τ*). As an example, we can imagine a simple system in which *z*_1_ the switch from non-leaky to leaky memory is activated only when the first input component *x*_1_ has high positive values. In the context of audition, it could correspond to a vocalization occurring in a given frequency band.

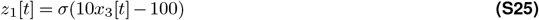

The memory of the *h*_1_ component of the hidden state of such a system only depends on the 3rd input element. With small values of the latter, the update gate is closed and the current form of *h*_1_ is maintained. However, with large values of *x*_3_, the update gate opens up and *h*_1_ is changed.

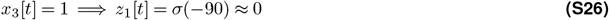

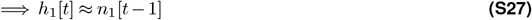

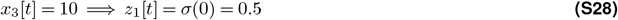

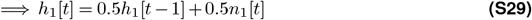

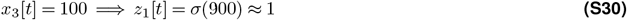

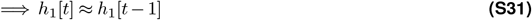

Note that this is just an example for illustration. In practice, much more complex patterns of inputs and hidden state are used to open and close the update gate. Thus, gated RNNs such as the GRU can dynamically encode, retain, and forget memory traces of input features in their hidden state over extended numbers of time-steps.

### Gain control from amplitude

We showed above that GRUs can dynamically remember or forget stored memory traces based on certain input patterns; one of these patterns could be the input amplitude, in whole or in part. It is indeed widely accepted that sensory neurons adapt their responses based on current or past stimulus amplitude, that is, mean stimulus level also affects neural gain (Rabinowitz et al. (2011)). In the previous example (Eq. S6-S12), the “key” to control the opening/closing of the update gate was a specific input component (*x*_3_). This behavior could be obtained by setting the weight of this component to a positive value, and the weight of other components to zero. But a positive weighted sum of all current input components of *x*[*t*] could serve as well as a measure of overall amplitude:

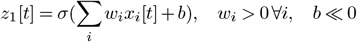

Note that most cochleagram representations of sound are positive, therefore ridding any potential problems due to negative inputs.

### Instantaneous contrast gain control

Another mechanism that has been extensively observed in sensory systems, and notably in the auditory pathway, is *contrast* gain control (Rabinowitz et al. (2011)). It is possible to approximate it from *amplitude* gain control (see above). Instead of using the global input amplitude as a “key”, we could rather take some measure of contrast in the current input vector *x*[*t*]. A valid definition –among others– for this could be the difference in amplitude between a target domain of the stimulus and the remaining “*background* “. Taking again the example of speech or animal vocalizations, auditory energy typically falls into a narrow range of relevant frequency bands: humans have for example peak sensitivity around 2,000 - 5,000 Hz corresponding to speech (Oxenham (2018)). With our proposed definition of contrast, the first component *h*_1_ of the hidden state could be updated depending on the following gate *z*_1_:

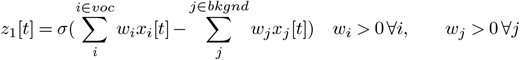

### Temporal correlation gain control

We showed that GRUs can compute and store (non-)leaky memory traces in their hidden state *h*, possibly over long sequences. In particular, we showed that gain control can be achieved through constant leak rates, or vary depending on the current stimulus amplitude, in whole or in parts, or on differences in amplitude between some of its parts (i.e., contrast). Sensory neurons can also modulate their responses as a function of the stimulus temporal statistics, like a temporal correlation (Natan et al. (2016)).

Let us define a very simple measure of temporal correlation, say in the first frequency band *x*_1_ of the stimulus *x*, as an exponential moving average (EMA) of the signal over a recent past, with more weight given to the most recent timesteps. With this definition, stable signals tend to give higher EMAs and noisy signals lower values.

A measure of temporal correlation can therefore be extracted by the GRU as the difference between the current value of the input and its leaky trace.

### Implementing AdapTrans with GRUs

AdapTrans is a recently proposed, general descriptive model of auditory ON and OFF responses and adaptation, that has been shown to consistently improve neural response fitting performances of stateless models (Rançon et al. (2024)). Applied within each frequency band of the cochleagram, it can be described essentially as computing the weighted difference between the current signal value and a EMA of the latter in a recent past. In other terms, it is an IIR filter that can be defined with the following recursive equation:

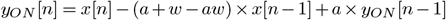

From the demonstrations provided above, we can show that GRUs can also approximate AdapTrans. Indeed, GRUs can compute an EMA of an input stimulus feature *x*_1_ with a certain time constant and store it into a hidden state element, for example *h*_2_. *z*_2_ can be chosen to reflect the exponential time constant (AdapTrans parameter *a*). At the current time-step *t*, this stored EMA value can be used to compute a weighted difference (AdapTrans parameter *w*) with the value of the current input *x*_1_[*t*]. AdapTrans is already a very general model of adaptation which encompasses prior approaches like IC adaptation (Willmore et al. (2016)). Thus, as GRUs are capable of approximating AdapTrans, they are capable of approximating these more specific models too.

### The strength that prior models of adaptation do not have

What essentially makes GRUs so powerful is their vector formulation, allowing them to compute and store both global and local features at the same time, for later use. These intermediate calculations enable very elaborate gain control mechanisms. In comparison, AdapTrans, IC adaptation and STP (Lopez Espejo et al. (2019)) rules only work frequency-wise and only *within* each spectral band. Prior studies have suggested that different frequency bands have indeed different adaptation parameters, but this alone might not be sufficient to truly reflect the behavior of actual neurons. Instead, we can see in our results that taking into account the contributions from neighboring and distant frequencies with RNNs in vector formulation, allows better fits to neuronal responses. This is consistent with the idea that neurons modulate their dynamics from signal statistics outside of their receptive fields (Rabinowitz et al. (2011)).

### Capabilities of other RNN models

We demonstrated that GRUs have the computational power to implement, or at least approximate, a wide variety of functions that have been extensively observed in biological systems, and notably in the auditory pathway. These demonstrations were performed on the GRU because this model leads to the best performances at the neural response fitting task in our study. Nonetheless, the same demonstrations hold for LSTMs because these models are also gated RNNs but with more equations and parameters and as such, can reproduce the functions performed by the GRUs.

On the other hand, the Elman (vanilla) RNN models are capable of implementing a constant –null or positive– leak, but quickly encounter some limitations, precisely because their leak is fixed and cannot depend on spectro-temporal stimulus statistics or the hidden state itself. As a reminder, the Elman RNNs are defined by:

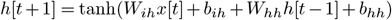

Therefore, contrast gain control or temporal correlation gain control are not achievable with these architectures, which could explain their lesser performances compared to the family of gated RNNs.

**Figure S3.**
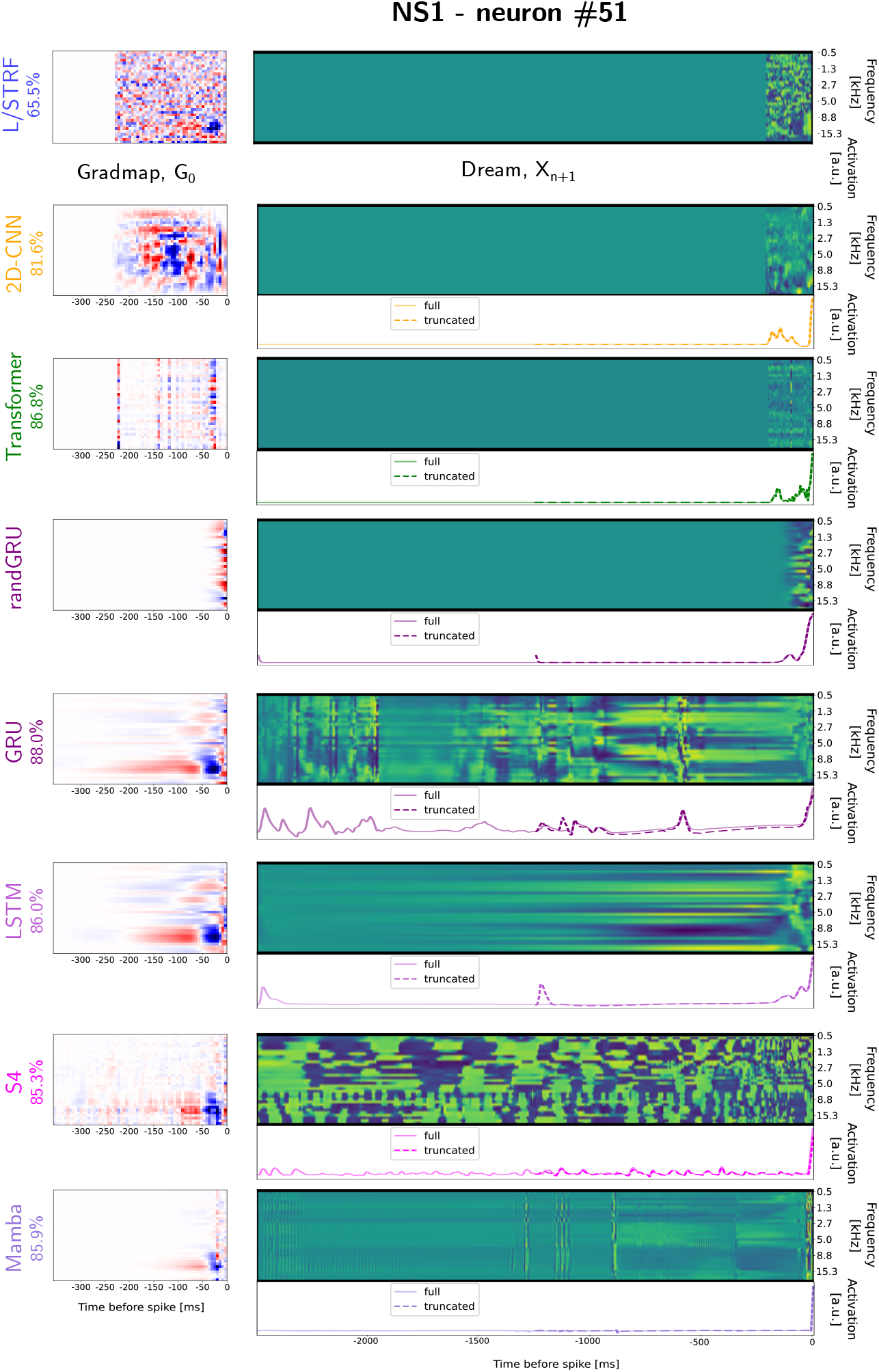
Gradmaps, extended dreams, and responses of other models for another NS1 neuron. *“randGRU”* : randomly initialized and untrained StateNet GRU model. left of each panel: gradmap *g*_0_. middle: dream *x*_1500_. bottom: predicted model response to the dream.

## Footnotes

1 https://crcns.org/data-sets/aa/aa-1/about

2 https://crcns.org/data-sets/ac/ac-1

